# Evaluating eDNA Detection from Live and Dead Control Sources

**DOI:** 10.64898/2026.04.27.721176

**Authors:** Scott M. Blankenship, Cheryl Dean, Katie Karpenko, Myfanwy Johnston, Matthew B. Espe, Gregg Schumer

## Abstract

Environmental DNA (eDNA) methods offer a powerful tool for monitoring aquatic species, yet field applications remain challenged by uncertainty in DNA transport, mixing, and detection, particularly in flowing or tidally influenced systems. One approach to improve confidence in eDNA surveys is the use of controlled DNA sources (positive controls), but questions remain regarding how the biological condition of the source influences eDNA release and detectability.

This study evaluated differences in eDNA concentrations emitted from live versus dead fish in a controlled, shallow, well-mixed channel. Using a fixed point-sampling design, we measured eDNA concentrations over time and modeled the effects of treatment, sampling time, temperature, and water velocity. Dead fish consistently released significantly higher concentrations of eDNA than live fish, while eDNA concentrations declined over time in both treatments. Water temperature and velocity did not significantly influence detection, and the rate of eDNA decline was similar between live and dead treatments.

These findings highlight the importance of source condition and site-specific mixing dynamics when interpreting positive control experiments and underscore the value of site characterization when designing eDNA sampling protocols.

## Introduction

Environmental DNA (eDNA) has revolutionized aquatic species monitoring by offering a sensitive, non-invasive method to detect the presence of target organisms in aquatic systems (Lawson Handley 2015). These approaches are particularly valuable for monitoring rare, cryptic, or protected species (Renan et al. 2017; Mauvisseau et al. 2019). Emerging methods for detection of eDNA have proven successful at monitoring biodiversity and do not require direct contact with organisms (Rees et al. 2014; Miya et al. 2020; Nagarajan et al. 2022).

The movement and dispersal of eDNA in flowing and tidally influenced waters remains poorly understood compared to unidirectional river systems. Factors such as current velocity, mixing patterns, and local hydrology can significantly affect the distribution of eDNA, leading to uncertainty about whether detected DNA accurately reflects the presence and/or concentration of target molecules (Pannone 2014). Without a clear accounting of how eDNA plumes disperse within complex systems, sampling designs risk both inaccurate characterization of DNA concentration from in situ collections and increased false negative rates from spatial mismatches between sampling points and eDNA sources (Dimian et al. 2013; Pannone 2014; Shogren et al. 2017). The dispersion of pollutant molecules has long been a focus of water quality control, which generally uses analytical models of mass transport and in situ measurement for validation of expected concentration distribution. While the behavior of DNA in the environment (persistence, shedding rates, transport) has been the subject of active investigation, the influence of water movement on sample collection itself has received less attention (Harrison et al. 2019; Blackman et al. 2024). Yet, the concentration of eDNA detected has been shown to vary due to transport dynamics (Laporte et al. 2020; Andruszkiewicz Allan et al. 2021; Thalinger et al. 2021), and velocity may interact with bathymetry to influence dispersion distance (Pont et al. 2018).

While eDNA detection has emerged as a transformative tool for monitoring aquatic species, several unresolved challenges limit its widespread application in ecosystems with complex hydrodynamics. These challenges arise from the dynamic and variable nature of eDNA transport in aquatic environments, as well as methodological uncertainties in detecting and interpreting eDNA detection data (Goldberg et al. 2016). Current eDNA sampling methodologies often lack robust validation mechanisms, making it difficult to assess their effectiveness in different environmental contexts. Incorporating known DNA sources (control DNA) into sampling designs has been proposed as a potential solution, but the utility and limitations of this approach remain underexplored in field applications (Jo et al. 2019). This study explores the utility of using control DNA sources—deliberately introduced DNA from known targets—to validate sampling protocols and adaptively refine detection methods. This work builds upon previous efforts evaluating eDNA methodologies, including the development of the {artemis} analysis framework (Espe et al. 2022), which provides robust statistical tools for modeling eDNA detection data.

The goal of this study was to increase the effectiveness of eDNA sampling as a tool for detecting aquatic species in complex ecosystems. To address challenges associated with detection reliability, we evaluated the use of control DNA sources—deliberately introduced DNA from a known quantity of organisms—to reduce uncertainty and enable adaptive refinement of sampling protocols. Preliminary analysis (not shown) of existing data on DNA concentration versus distance from control sources and water volume collected led us to focus on a key unresolved question: does the concentration of DNA ([DNA]) emitted differ between live and dead DNA sources? This is important because many studies rely on frozen tissue or preserved material as proxies for live organisms, yet the degree to which these mimic natural eDNA shedding remains unclear. We hypothesized that (1) there is a measurable difference in eDNA concentration shed by live and dead fish at 20-minute intervals over a 45-minute period post-deployment, and (2) live and dead fish exhibit different eDNA shedding dynamics over time.

## Materials and Methods

### Experimental Design

To test hypotheses related to differences in DNA detections from live and dead DNA sources, we designed a complete block experiment where the treatment of interest was live fish or a comparable biomass of dead, previously frozen fish of the same species. Comet goldfish (*Carassius auratus*) were used for the DNA source. The fish came in two distinct size groups: small (1.2-4 cm total length) and large (∼5-7.5 cm total length). There were 120 small and 80 large fish available for the experiment. Within size groups, half the fish were randomly assigned to the dead treatment and euthanized according to a standard accepted protocol (overdose of MS-222; Jenkins et al. 2014). The dead fish were immediately frozen for at least 10 hours prior to being used in an experimental block. Live fish were held and cared for in ∼151-Liter oxygenated tanks before being randomly selected (within size groups to maintain comparable biomasses across blocks) for blocks on the day(s) of the experiment.

The experiment was conducted in Newtown Ditch in Nevada City, California. This is an idealized, narrow (2 m-3 m wide), shallow (0.33 m-0.65 m) channel of water with unidirectional flow with no known occurrence of *C. auratus*. Cages containing live or dead fish (live/dead cars) were deployed 50 m upstream of the DNA sampling point at an average depth of 0.26 m. The 50 m stretch of experimental channel was chosen because it included a short culvert with a shallow drop-off ∼30 m downstream of the fixed live/dead car location that contributed to DNA particle mixing relative to points taken upstream of it. Thus, we assumed the eDNA from the cage location was uniformly distributed in the water column at the fixed DNA sampling location.

Each sampling block, consisting of deployments of both live and dead fish treatments, was replicated five times over two days. We disclose that in the field, the live/dead treatment deployment sequence was not randomized. Rather, for each block the deployment was live fish treatment followed by dead fish treatment. Between each fish treatment within blocks, after the live fish treatment cage was removed, a period of 30 minutes elapsed without any treatment deployed to allow for the clearing of eDNA from the system. Results need to be interpreted accordingly, given the inadvertent de-randomization of live/dead treatments does not achieve the ideal complete block experiment design conditions.

Control samples were taken following the clearing period (post live fish treatment) and before the deployment of the next dead fish treatment to verify the eDNA cleared from the system. We assumed the clearing time of 30 minutes was sufficient such that deployment of one treatment did not affect subsequent deployments. To account for environmental difference between and within days, water temperature and velocity were recorded when each treatment was deployed.

#### eDNA Collection Methods

Environmental DNA was collected during this study using point sampling. Water samples of 1000 mL volume were taken 5, 25, and 45 minutes after the treatment (live car or dead car) began. Each sample was filtered through a Sterivex™ (0.45 um PVDF membrane) (MilliporeSigma) eDNA filter. Pumps were affixed in parallel to allow for simultaneous filtration of three filter replicates and powered using a brushless, cordless drill with a 12.7 mm spade bit attachment. The filtrate from each sample was captured in a graduated cylinder and the volume filtered was recorded along with the date and event identifier. When filtration ended, water was removed from the filter cavity and the filter was capped on both ends using VWR nylon male and female luer lock caps. Samples were placed into sterile Ziploc bags and stored on ice for the duration of the field day, then transferred to a −20° C freezer at the end of the day to stabilize eDNA. We used sterile gloves at each event to reduce the risk of cross-contamination and collected field controls at the start of each field day to confirm sample integrity. Field negative controls were collected by filtering 500 mL of ultrapure water as described above.

### Laboratory Protocols

#### Environmental DNA Laboratory Analysis

DNA was isolated from each Sterivex™ eDNA filter following a modified (Miya et al. 2016) protocol. The presence of target DNA was determined by using a species-specific qPCR assay for Comet Goldfish (Kamoroff and Goldberg 2018), with three technical qPCR replicates per filter eDNA extraction. The qPCR amplification mixture final volume of 10 ul included 4 ul template, 5 ul 2x Applied Biosystems TaqMan Environmental PCR Master Mix and varying final primer and probe concentrations. Thermocycling was performed using the Applied Biosystems QuantStudio™ 3 following: 10 min at 95° C, then 40 cycles of 15 sec at 95° C and 1 min at assay specific annealing temperature. Each qPCR plate included three no template controls (ultra-pure water) and two positive controls (2 ng/µl Comet Goldfish and Ballyhoo DNA). All PCR master mixes were made inside a UV sterilized PCR enclosed workstation. All PCR reactions were conducted on instruments located outside of the main lab in a separate portion of the building. Results of the qPCR reactions were analyzed using QuantStudio™ Design & Analysis Software with the magnitude of the qPCR signal reported as quantification cycle (Cq).

### 2.3 Analytical Framework

The {artemis} R package (v 2.0.0, Espe et al, 2022) in the R language (v 4.1.3, R Core Team, 2022) was utilized to analyze eDNA data from the controlled experiment. The artemis package was developed specifically to model qPCR data from eDNA experiments and fully accounts for uncertainty introduced by data censorship inherent in the qPCR process. This analysis framework allowed modeling of the relationship between DNA concentration and environmental variables, assessing the sensitivity and specificity of eDNA detection under varying conditions, and comparing detection probabilities between controlled and field conditions. The treatment (live or dead), timepoint, and the interaction between treatment and timepoint were modeled as fixed effects, with block modeled as a random effect. The covariates of water temperature (degrees Celsius) and water velocity (meters per second) were also included in the model to account for systematic differences among blocks.

## Results

### Live versus Dead DNA Sources

During the experiment, water temperature ranged from 11.9° to 15.0° C, while water velocity ranged from 0.56 to 0.84 meters per second. Both covariates were found to be uncorrelated to treatment deployment (analysis not presented). The DNA detections (Cq) by filter and technical replicate were provided in supplemental data. The field controls and laboratory process controls were all negative for DNA detections (supplemental data).

Environmental DNA concentrations differed consistently between live and dead fish treatments throughout the experiment. Raw Cq values showed lower Cq (higher eDNA concentration) in samples associated with the dead fish treatment relative to the live fish treatment across all blocks and timepoints (Figure 1). In both treatments, Cq values increased through time, indicating declining eDNA concentration following deployment.

**Figure 1.**
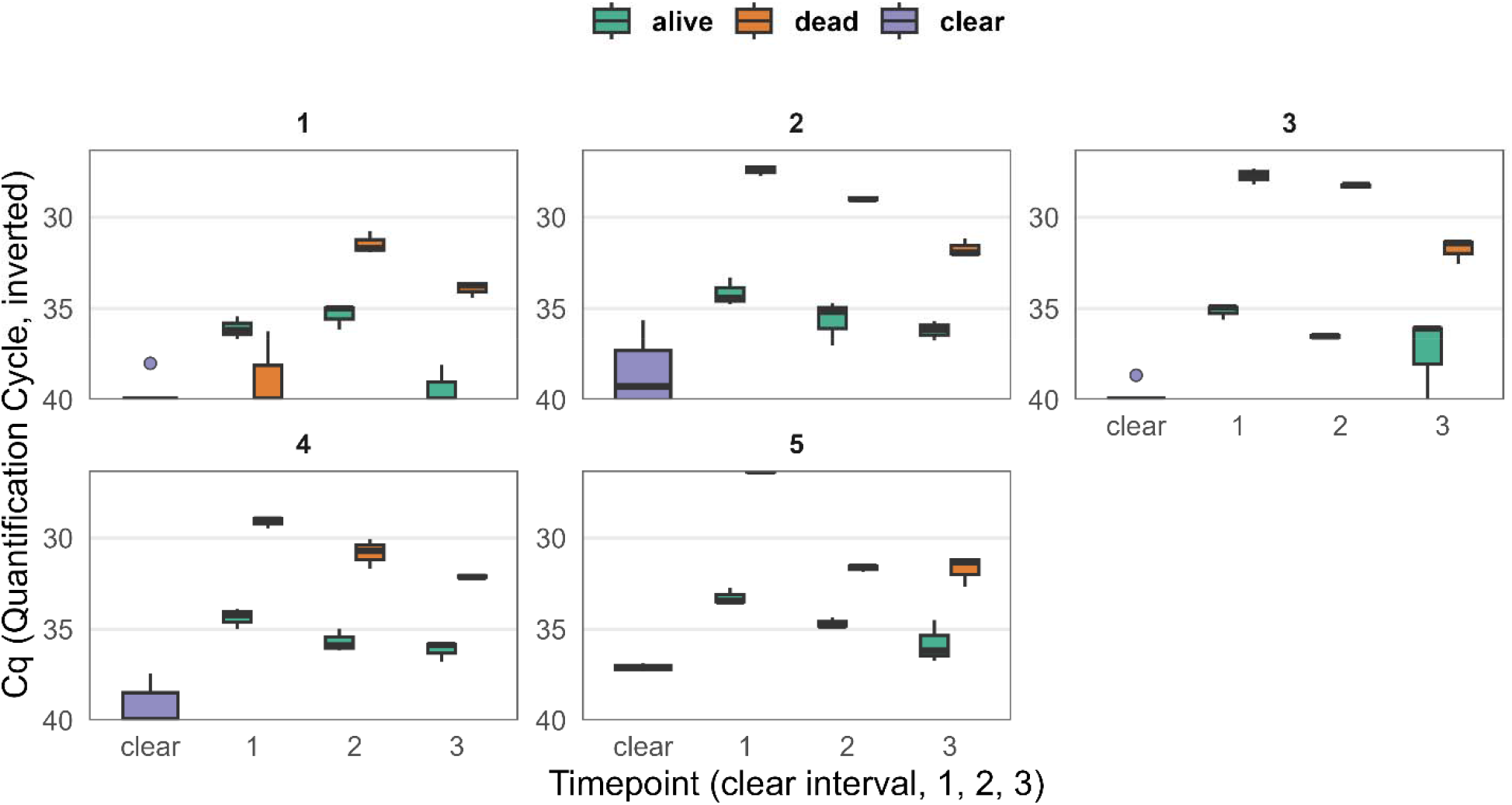
Quantification cycle (Cq) values for Target = GOLD eDNA across experimental blocks, treatments, and timepoints. Boxplots show the distribution of technical replicates for each treatment (alive, dead, clear control) within each block (panels 1–5) and timepoint (clear interval, 1, 2, 3). Lower Cq values indicate higher eDNA concentration; the y-axis is inverted for interpretability.

Following statistical modelling of DNA detections, the estimated effect sizes for the live versus dead eDNA experiment are presented in Figure 2. Note that if a parameter value does not overlap with zero, then the observed effect (either negative or positive) was statistically significant. Positive values indicate higher eDNA concentration. A regression model describing ln-transformed eDNA concentration supported the patterns observed from Cq data. Biomass was included to control for differences in total fish mass among blocks and treatments and to estimate treatment effects on eDNA shedding independent of fish size. After accounting for biomass, the dead fish treatment was associated with a strong positive effect on eDNA concentration relative to the live fish treatment (mean 3.8; std. dev. 0.58), while time since deployment showed a consistent negative effect (mean −0.8; std. dev. 0.30), indicating declining eDNA concentration over time. Water temperature and water velocity exhibited weak or uncertain effects under the range of conditions observed. The interaction between treatment and time was not supported in the updated model, suggesting that the rate of eDNA decline through time was similar for live and dead fish treatments despite differences in initial concentration. The “sigma” estimate was (std. dev. 0.22). This value represented the standard deviation of the model’s residuals, suggesting that treatment, time, temperature, and velocity factors do not account for all the variability in eDNA detections.

**Figure 2.**
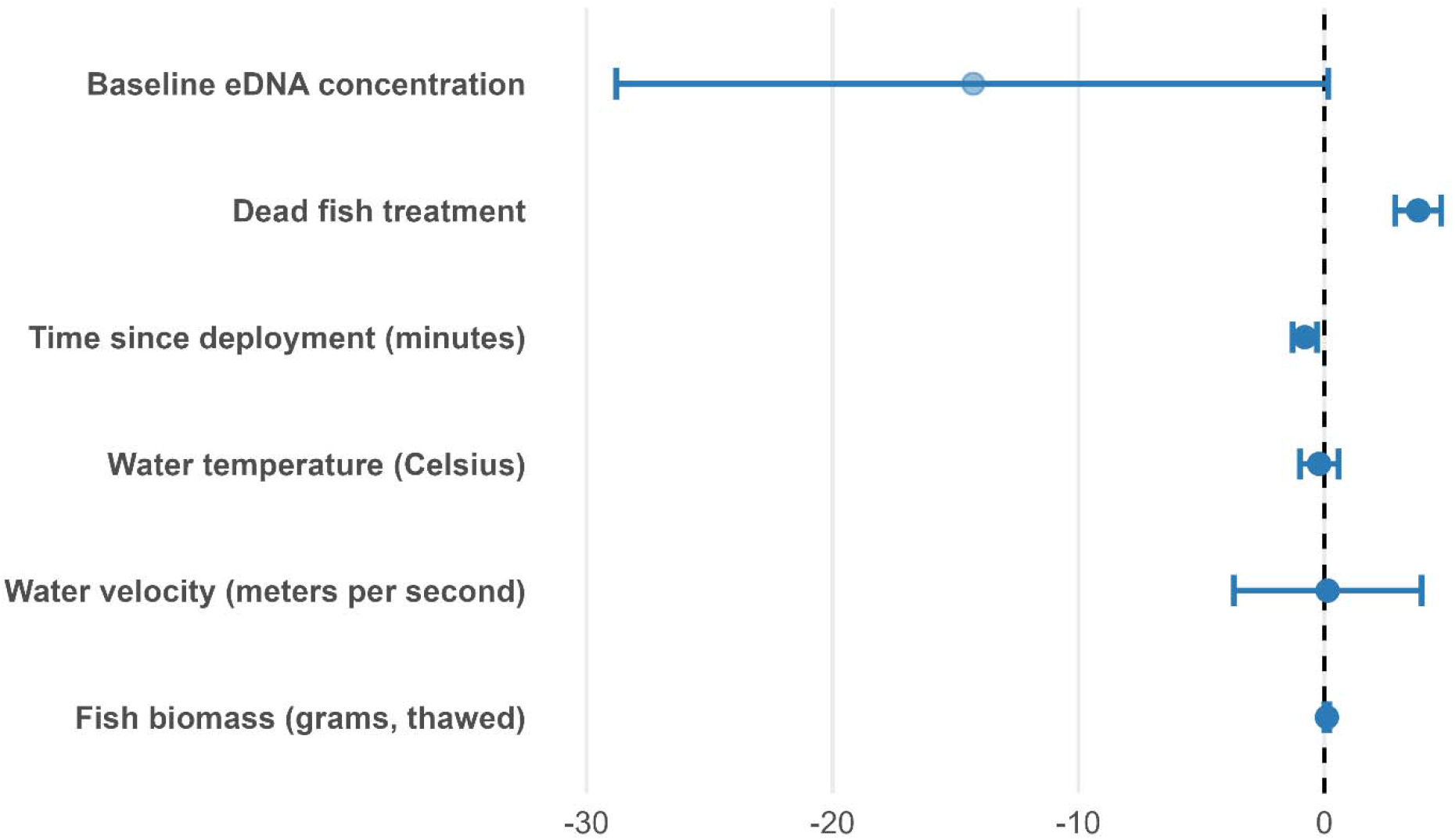
Estimated effects (posterior means ± 90% credible intervals) from a regression model describing variation in ln-transformed eDNA concentration during a live versus dead fish experiment. Positive values indicate higher eDNA concentrations, while negative values indicate lower concentrations relative to the reference condition.

To assess whether differences in eDNA concentration reflected differences in biomass-normalized shedding, eDNA concentrations were converted to standard-curve–derived concentration equivalents and normalized by total fish biomass deployed in each block. Biomass-normalized eDNA concentrations were consistently higher for the dead fish treatment than for the live fish treatment across all blocks and timepoints (Figure 3). Although absolute concentrations declined through time in both treatments, the relative separation between treatments remained evident, indicating similar temporal decay patterns. Corroborating this observation was that the interaction term between treatment and timepoint was not significant in regression model. Within each block, technical replicates showed variability relative to the magnitude of the treatment effect (Figure 3), and the overall pattern of higher biomass-normalized eDNA shedding from dead fish was consistent across blocks. Block 5 was excluded from biomass-normalized analyses due to missing biomass measurement for the live fish treatment.

**Figure 3.**
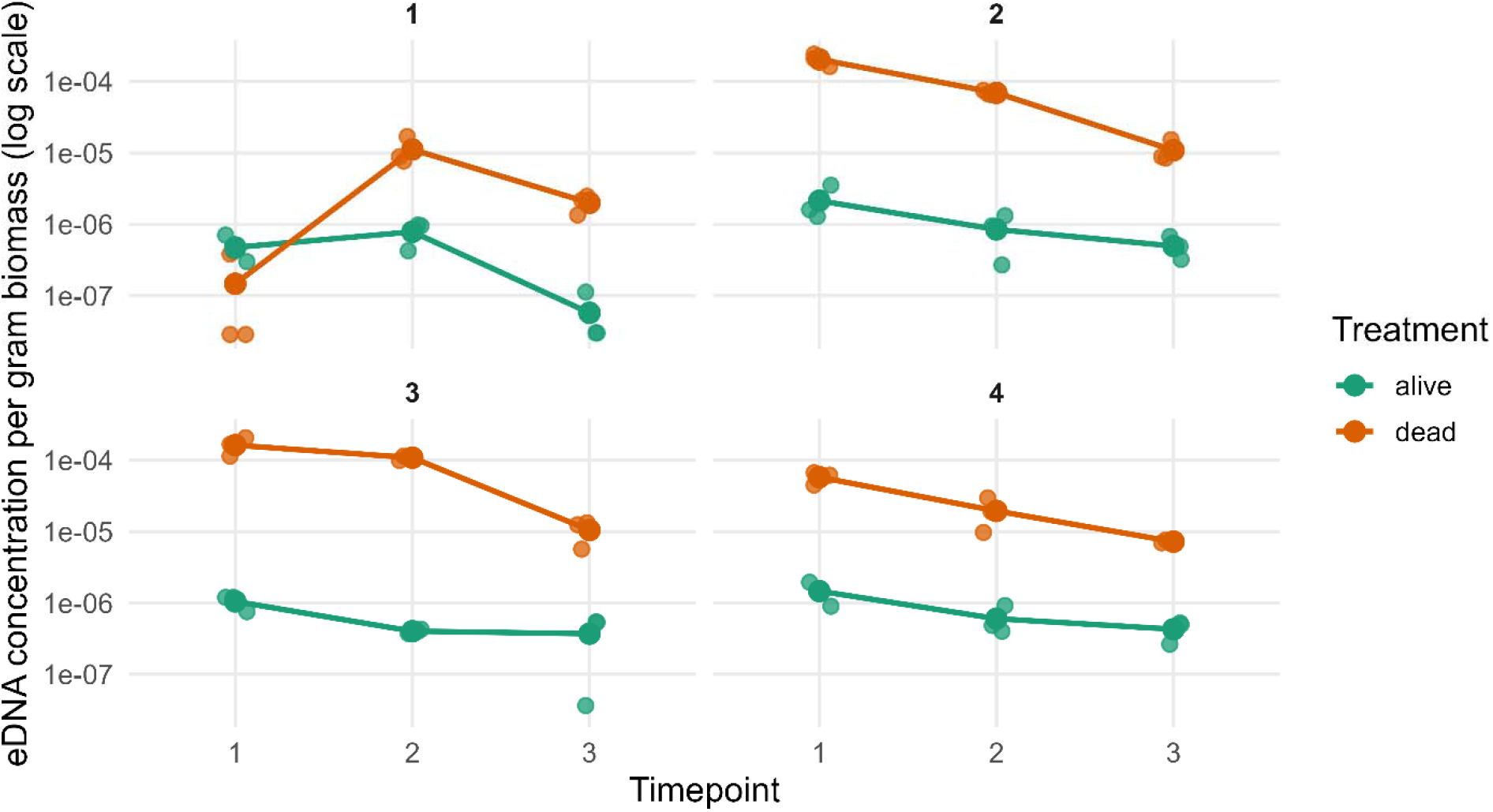
Biomass-normalized eDNA concentration for live and dead fish treatments (Target = GOLD). Points represent individual technical replicates, and lines show mean eDNA concentration per gram biomass at each timepoint. Values are shown on a log scale. Blocks 1–4 are shown; Block 5 was excluded due to missing biomass measurement.

## Discussion

### Live versus Dead DNA Sources

This study found that dead fish released significantly more eDNA than live fish, and that eDNA concentrations declined consistently over time for both source types (Figure 1). After accounting for fish biomass and environmental covariates, DNA source condition (live versus dead) and time since deployment emerged as the dominant predictors of eDNA concentration in this controlled, well-mixed channel (Figure 2). In contrast, water temperature and velocity showed no detectable effects within the observed experimental range. These results provide a quantitative foundation for understanding how biological state influences eDNA availability in flowing environments under conditions of effective mixing.

While the regression results isolate source condition and time as primary drivers under controlled conditions, interpretation of live and dead eDNA sources requires consideration of how DNA disperses from the source. A key assumption in many controlled eDNA studies – particularly those using enclosed organisms (live cars or dead cars) – is that DNA disperses uniformly from the source. However, prior work and our own observations challenge this assumption. Live cars introduce a concentrated plume of DNA into the system, but the rate of mixing is governed by site-specific hydrodynamics, resulting in spatially variable eDNA (Shogren et al. 2017). When dispersal is relatively uniform, as was in the present experiment, variability in eDNA detection is lower and treatment effects are more readily observed. In contrast, when mixing is incomplete— as is often the case in field deployments at sites with complex or unknown hydrology—sampling location becomes a major source of uncertainty. This may result in lower-than-expected detections near the source or a weak relationship between sample volume and [DNA]. These effects may not reflect underlying biological or methodological error, but rather hydrodynamic artifacts (Deiner and Altermatt 2014; Shogren et al. 2017). Spatially structured transect sampling has been used in a variety of systems to better characterize DNA transport and dispersal, and can help mitigate the effects of uneven mixing by providing a more comprehensive picture of eDNA distribution across a site (Jerde et al. 2016; Pont et al. 2018). However, even transect-based approaches are sensitive to flow conditions and local bathymetry, and uniformity remains a critical factor when interpreting eDNA monitoring data.

### Interpreting Source Differences

The observed difference in eDNA concentrations between live and dead sources has important implications for calibration experiments. Although dead fish consistently produced higher eDNA concentrations, the rate of decline through time was similar between treatments, suggesting that source condition primarily affects initial shedding magnitude rather than short-term decay dynamics. The experiment was not designed to resolve fine-scale degradation processes, limiting our ability to distinguish between treatment-specific differences in eDNA decay and physical dilution or transport over longer time scales. Other studies have found that dead fish shed large amounts of eDNA initially, followed by rapid tapering (Maruyama et al. 2014; Jo et al. 2019), whereas live fish may eventually reach a steady-state shedding rate. Importantly, treatment effects persisted after normalizing eDNA concentrations by thawed fish biomass, indicating that dead fish shed more eDNA per unit mass rather than simply reflecting differences in total tissue. Additionally, we observed evidence of eDNA “pulsing” following handling or jostling (as seen in Ballyhoo dead car data, not presented), which may or may not reflect natural shedding in free-swimming organisms (Tillotson et al. 2018). Finally, because the experiment was conducted in a narrow, shallow, and well-mixed channel, it may not be directly generalized to estuarine or stratified systems. Future validation studies should account for hydrologic differences at sites with dynamic (tidally influenced), variable (seasonal), or unknown (confluences, lacustrine, manmade structures) hydrology when designing both live and dead car eDNA trials.

### Methodological Insights

While the biological patterns observed here were robust, absolute eDNA concentrations necessarily depend on the specific filtration hardware, laboratory protocols, and analytical choices used. While we have neither altered our filtration hardware nor changed laboratory protocol in this experiment, ongoing work continues to evaluate how both field and lab methodological choices influence eDNA capture. Previous work indicates that methodological differences can substantially affect detection probability and estimated concentrations, although such effects are typically smaller than the treatment-level differences observed in this study (Goldberg et al. 2016; Wilcox et al. 2018). The following recommendations reflect both the findings of this study and our broader experience working in dynamic aquatic systems. These are intended to maximize the detection probability of positive controls and are best interpreted as a living document, evolving with future data and insights.

#### Recommendations for Live car and Dead car Experiments

1. Control species should be absent from the study system to avoid confounding detections. Note that the bait species (e.g., anchovy) are not guaranteed to be absent, especially in estuarine or recreational areas.
2. Dead fish released higher eDNA concentrations per unit biomass than live fish in this study, as demonstrated by both regression analyses and biomass-normalized comparisons. To maximize consistency in shedding dynamics when using dead fish as positive controls, freshly frozen individuals that have not undergone repeated freeze– thaw cycles should be used, as repeated thawing may alter tissue integrity and eDNA release patterns.
3. Detection probabilities decrease over time for both live and dead sources. Highest detections typically occur within the first 0–2 hours post-deployment. Longer deployment times increase variability in [DNA].
4. Dead fish are expected to shed more eDNA than live fish within the first 0–2 hours post-deployment, reflecting higher initial shedding. eDNA concentrations declined through time in both treatments, but fine-scale temporal shedding dynamics were not resolved in this study. For consistency, the same control species should be used across experiments.
5. To minimize variability and isolate covariates affecting detection probability in tidally influenced systems, sampling should occur during periods of unidirectional flow (outgoing or incoming tides). Consistent flow direction helps maintain a stable eDNA plume orientation, improving comparability among samples and reducing variability associated with changing hydrodynamic conditions.
6. Sampling points or transects should intersect the eDNA plume where it is most homogenously mixed to minimize spatial artifacts.
7. Water volume filtered, distance from the source, and biomass of the positive control should be scaled to site-specific hydrodynamics, such as flow directionality (unidirectional vs. reversing), flow velocity, turbulence and eddying, and channel geometry, as well as the desired spatial detection range.

## Conclusions and Future Directions

Environmental DNA has emerged as a promising tool for aquatic monitoring, yet methodological uncertainties still constrain its utility in complex systems. This study demonstrates that source condition (live or dead) can strongly influence eDNA concentrations, and that time remains a critical factor in detectability. These findings highlight the need for standardized positive control protocols, control sources, and better site characterization in field-based eDNA studies. Future efforts should explore the behavior of positive control eDNA across a wider range of hydrologic conditions, enclosure designs, and species types. Improved characterization of study sites – particularly flow regime, mixing behavior, and stratification – can help refine sampling designs and detection models. Integrating spatially resolved transect sampling with real-time hydrodynamics monitoring could better capture eDNA plume behavior and improve predictability. An unresolved question for eDNA interpretation concerns the persistence and downstream detectability of eDNA originating from dead organisms. While decay rates of free-floating eDNA are increasingly well characterized, less is known about how long biomass-associated eDNA from carcasses remains detectable under different hydrologic conditions. Inference about whether detected eDNA reflects recent biological presence or legacy sources is complicated, particularly when based on isolated or single-point detections. Ultimately, a flexible, system-specific approach to validation using positive controls will enhance the reliability of eDNA applications for management and conservation, particularly when live or dead organism controls are used to interpret detection probabilities in flowing systems.

## Acknowledgements

We would like to acknowledge Jasmine Williamshen and Michelle Leung for their diligent assistance with implementing field experiments. Special thanks to Chip Close (Nevada Irrigation District) for allowing the use of their water conveyance facility for experimentation. Special thanks to Shawn Acuña (Principal Resource Specialist; Metropolitan Water District) for helpful comments on manuscript and DNA detection applications within the regulatory environment.

Funding for this study was provided by the State Water Contractors, (contract no. 21-29).

Fish transport was completed in oxygenated hauling tanks, with dissolved oxygen levels and temperature being checked at set intervals during transport to ensure optimal conditions for each species. After each experiment, all fish were euthanized with tricaine methane sulfonate (Syndel USA Tricaine-S, MS-222) in accordance with permit specifications and professional society guidelines (Jenkins et al. 2014). Fish handling, transport and euthanasia followed specifications outlined in the listed permits, agreements, and Fisheries Society guidelines.

The views expressed are those of the authors and do not represent the official opinion of any employer, institution, or government agency. The authors declare that they have no known competing financial interests or personal relationships that could have appeared to influence the work reported in this paper.

## Authors Contributions

SB made major to the interpretation of the data and writing of the manuscript. CD made major contributions to the acquisition and analysis of the data, KK, ME, MJ made major contributions to the design of the study, acquisition, analysis, and interpretation of the data, and writing of the manuscript. GS made major contributions to the conception of the study and writing of the manuscript.

## Data Availability

The data that supports the findings of this study are available on request from the corresponding author Dr. Scott Blankenship.

## Ethics in Publishing Statement

I testify on behalf of all co-authors that our article submitted followed ethical principles in publishing.

Title: Evaluating eDNA Detection from Live and Dead Control Sources

All authors agree that:

This research presents an accurate account of the work performed, all data presented are accurate and methodologies detailed enough to permit others to replicate the work.

This manuscript represents entirely original works and/or if work and/or words of others have been used, that this has been appropriately cited or quoted and permission has been obtained where necessary.

This material has not been published on the whole or in part elsewhere.

The manuscript is not currently being considered for publication in another journal.

That generative AI and AI-assisted technologies have not been utilized in the writing process or if used, disclosed in the manuscript the use of AI and AI-assisted technologies and a statement will appear in the published work.

That generative AI and AI-assisted technologies have not been used to create or alter images unless specifically used as part of the research design where such use must be described in a reproducible manner in the methods section.

All authors have been personally and actively involved in substantive work leading to the manuscript and will hold themselves jointly and individually responsible for its content.

## References

Andruszkiewicz Allan E, Zhang WG, C. Lavery A, F. Govindarajan A. 2021. Environmental DNA shedding and decay rates from diverse animal forms and thermal regimes. Environ DNA. 3:492–514. 10.1002/edn3.141

Blackman RC, Carraro L, Keck F, Altermatt F. 2024. Measuring the state of aquatic environments using eDNA— upscaling spatial resolution of biotic indices. Philos Trans R Soc B Biol Sci. 379:20230121. 10.1098/rstb.2023.0121

Deiner K, Altermatt F. 2014. Transport distance of invertebrate environmental DNA in a natural river. PloS One. 9:e88786.

Dimian MFA, Wadi AS, Ibrahim FN. 2013. The effect of added pollutant along a river on the pollutant concentration described by one-dimensional advection diffusion. Int J Eng Sci Technol. 5:1662.

Espe MB, Johnston M, Blankenship SM, Dean CA, Bowen MD, Schultz A, Schumer G. 2022. The artemis package for environmental DNA analysis in R. Environ DNA. [accessed 2022 March 11]. 10.1002/edn3.277

Goldberg CS, Turner CR, Deiner K, Klymus KE, Thomsen PF, Murphy MA, Spear SF, McKee A, Oyler-McCance SJ, Cornman RS, Laramie MB, Mahon AR, Lance RF, Pilliod DS, Strickler KM, Waits LP, Fremier AK, Takahara T, Herder JE, Taberlet P. 2016. Critical considerations for the application of environmental DNA methods to detect aquatic species. Methods Ecol Evol. 7:1299–1307. 10.1111/2041-210X.12595

Harrison JB, Sunday JM, Rogers SM. 2019. Predicting the fate of eDNA in the environment and implications for studying biodiversity. Proc R Soc B Biol Sci. 286:20191409. 10.1098/rspb.2019.1409

Jenkins JA, Bart Jr HL, Bowker JD, Bowser PR, MacMillan JR, Nickum JG, Rose JD, Sorensen PW, Whitledge GW, Rachlin JW. 2014. Guidelines for the use of fishes in research. Bethesda Md USA Am Fish Soc. [accessed 2026 January 20]. [accessed 2026 Jan 20]. Available from: https://nnmc.edu/_document_repository/iacuc-docs/Guidelines-for-Use-of-Fishes.pdf

Jerde CL, Olds BP, Shogren AJ, Andruszkiewicz EA, Mahon AR, Bolster D, Tank JL. 2016. Influence of Stream Bottom Substrate on Retention and Transport of Vertebrate Environmental DNA. Environ Sci Technol. 50:8770– 8779. 10.1021/acs.est.6b01761

Jo T, Murakami H, Yamamoto S, Masuda R, Minamoto T. 2019. Effect of water temperature and fish biomass on environmental DNA shedding, degradation, and size distribution. Ecol Evol. 9:1135–1146. 10.1002/ece3.4802

Kamoroff C, Goldberg CS. 2018. An issue of life or death: using eDNA to detect viable individuals in wilderness restoration. Freshw Sci. 37:685–696. 10.1086/699203

Laporte M, Bougas B, Côté G, Champoux O, Paradis Y, Morin J, Bernatchez L. 2020. Caged fish experiment and hydrodynamic bidimensional modeling highlight the importance to consider 2D dispersion in fluvial environmental DNA studies. Environ DNA. 2:362–372. 10.1002/edn3.88

Lawson Handley L. 2015. How will the ‘molecular revolution’ contribute to biological recording? Biol J Linn Soc. 115:750–766. 10.1111/bij.12516

Maruyama A, Nakamura K, Yamanaka H, Kondoh M, Minamoto T. 2014. The Release Rate of Environmental DNA from Juvenile and Adult Fish. PLOS ONE. 9:e114639. 10.1371/journal.pone.0114639

Mauvisseau Q, Davy-Bowker J, Bulling M, Brys R, Neyrinck S, Troth C, Sweet M. 2019. Combining ddPCR and environmental DNA to improve detection capabilities of a critically endangered freshwater invertebrate. Sci Rep. 9:14064. 10.1038/s41598-019-50571-9

Miya M, Gotoh RO, Sado T. 2020. MiFish metabarcoding: a high-throughput approach for simultaneous detection of multiple fish species from environmental DNA and other samples. Fish Sci. 86:939–970. 10.1007/s12562-020-01461-x

Miya M, Minamoto T, Yamanaka H, Oka S, Sato K, Yamamoto S, Sado T, Doi H. 2016. Use of a filter cartridge for filtration of water samples and extraction of environmental DNA. JoVE J Vis Exp. e54741.

Nagarajan RP, Bedwell M, Holmes AE, Sanches T, Acuña S, Baerwald M, Barnes MA, Blankenship S, Connon RE, Deiner K. 2022. Environmental DNA methods for ecological monitoring and biodiversity assessment in estuaries. Estuaries Coasts. 45:2254–2273.

Pannone M. 2014. Predictability of tracer dilution in large open channel flows: Analytical solution for the coefficient of variation of the depth-averaged concentration. Water Resour Res. 50:2617–2635. 10.1002/2013WR013986

Pont D, Rocle M, Valentini A, Civade R, Jean P, Maire A, Roset N, Schabuss M, Zornig H, Dejean T. 2018. Environmental DNA reveals quantitative patterns of fish biodiversity in large rivers despite its downstream transportation. Sci Rep. 8:10361. 10.1038/s41598-018-28424-8

Rees HC, Maddison BC, Middleditch DJ, Patmore JRM, Gough KC. 2014. REVIEW: The detection of aquatic animal species using environmental DNA – a review of eDNA as a survey tool in ecology. J Appl Ecol. 51:1450– 1459. 10.1111/1365-2664.12306

Renan S, Gafny S, Perl RGB, Roll U, Malka Y, Vences M, Geffen E. 2017. Living quarters of a living fossil—Uncovering the current distribution pattern of the rediscovered Hula painted frog (Latonia nigriventer) using environmental DNA. Mol Ecol. 26:6801–6812. 10.1111/mec.14420

Shogren AJ, Tank JL, Andruszkiewicz E, Olds B, Mahon AR, Jerde CL, Bolster D. 2017. Controls on eDNA movement in streams: Transport, retention, and resuspension. Sci Rep. 7:5065.

Thalinger B, Kirschner D, Pütz Y, Moritz C, Schwarzenberger R, Wanzenböck J, Traugott M. 2021. Lateral and longitudinal fish environmental DNA distribution in dynamic riverine habitats. Environ DNA. 3:305–318. 10.1002/edn3.171

Tillotson MD, Kelly RP, Duda JJ, Hoy M, Kralj J, Quinn TP. 2018. Concentrations of environmental DNA (eDNA) reflect spawning salmon abundance at fine spatial and temporal scales. Biol Conserv. 220:1–11. 10.1016/j.biocon.2018.01.030

Wilcox TM, Carim KJ, Young MK, McKelvey KS, Franklin TW, Schwartz MK. 2018. Comment: The Importance of Sound Methodology in Environmental DNA Sampling. North Am J Fish Manag. 38:592–596. 10.1002/nafm.10055

